# MeCP2 represses the induction and maintenance of long-term synaptic plasticity

**DOI:** 10.1101/2022.10.25.513721

**Authors:** Rong-Yu Liu, Yili Zhang, Paul Smolen, John H. Byrne

## Abstract

Mechanisms of specific memory deficits associated with Rett syndrome are poorly understood, at least in part because mutations of *MECP2* have confounding effects on nervous system development and basal synaptic transmission. To mitigate such empirical uncertainties, this study exploited technical advantages of the *Aplysia* sensorimotor synapse to examine the potential role of MeCP2 in long-term synaptic plasticity. The results indicate MeCP2 may act as an inhibitory constraint on gene expression required for formation as well as maintenance of plasticity.

## Main

Rett syndrome, a neurodevelopmental disorder associated with memory deficits, is caused in the majority of cases (96%) by mutations of the X-chromosome linked gene *MECP2* ^1, 2^. MeCP2 acts, most commonly, as a transcriptional repressor when bound to promoters ^3^. Mouse genetic models revealed severe neurological abnormalities, including malformed dendrites, axons, and synaptic structures^4^, and deficits in synaptogenesis^5, 6^, and in learning and memory, with aberrant MeCP2 levels possibly altering long-term potentiation (LTP)^7-9^. Despite these advances, a specific role for MeCP2 in memory mechanisms is poorly understood. This incomplete understanding results, at least in part, from confounding effects of MeCP2 on basal synaptic transmission and short-term synaptic plasticity (e.g., paired pulse facilitation)^7, 10-12^, which might be expected to affect induction of LTP. Moreover, manipulations of MeCP2 to date affect both presynaptic and postsynaptic neurons. To reduce these complexities, the present study exploited the technical advantages of the *Aplysia* sensorimotor synapse to examine the potential role of MeCP2 in long-term synaptic facilitation (LTF), a well-established model system that has provided insights into general mechanisms of synaptic plasticity (see ^13^ for review). In this system long-term synaptic plasticity can be studied *in vitro* with the ability to separately monitor and control the presynaptic and postsynaptic neurons.

Three temporally distinct protocols have been used to induce LTF^14^, the Standard protocol consisting of 5 regularly spaced pulses of 5-HT (5P,S), the Enhanced protocol of 5 irregularly spaced pulses (5P,E) which induces stronger and more persistent LTF, and the 2-pulse protocol (2P). We validated specificity of a vertebrate MeCP2 antibody (αMeCP2) for MeCP2 in *Aplysia* neuronal ganglia (Online Methods and Supplementary Information, Fig. S1), and used αMeCP2 to examine changes in MeCP2 levels after applying these protocols to sensorimotor synapses. Compared to vehicle-treated control (n = 9), immunoreactivity to MeCP2 was decreased by all three protocols (n = 7 for 5P,S; n = 6 for 5P,E; n=5 for 2P) (Fig. 1A). One-way repeated measurement ANOVA indicated significant overall differences among the four groups (F_3,15_ = 47.23, p < 0.001). Student-Newman-Keuls (SNK) post hoc analysis indicated the MeCP2 level in Veh group was significantly greater than those in all three 5-HT treated groups (Veh *vs*. 5P,S: q = 11.33, p < 0.001; Veh *vs*. 5P,E: q = 15.93, p < 0.001; and Veh *vs*. 2P: q = 7.25, p < 0.001). There was also a significant difference in MeCP2 levels between the strong LTF-inducing protocol (5P,E) and the other protocols (5P,S and 2P) (5P,E *vs*. 5P,S: q = 4.91, p = 0.044; and 5P,E *vs*. 2P group: q = 6.81, p < 0.001). However, there was not a significant difference between 5P,S and 2P groups (q = 2.49, p = 0.099). These results indicate MeCP2 levels are downregulated by stimuli that induce LTF. Moreover, the extent of downregulation of MeCP2 correlates with the effectiveness of the protocols in inducing LTF ^15^.

**Figure 1.**
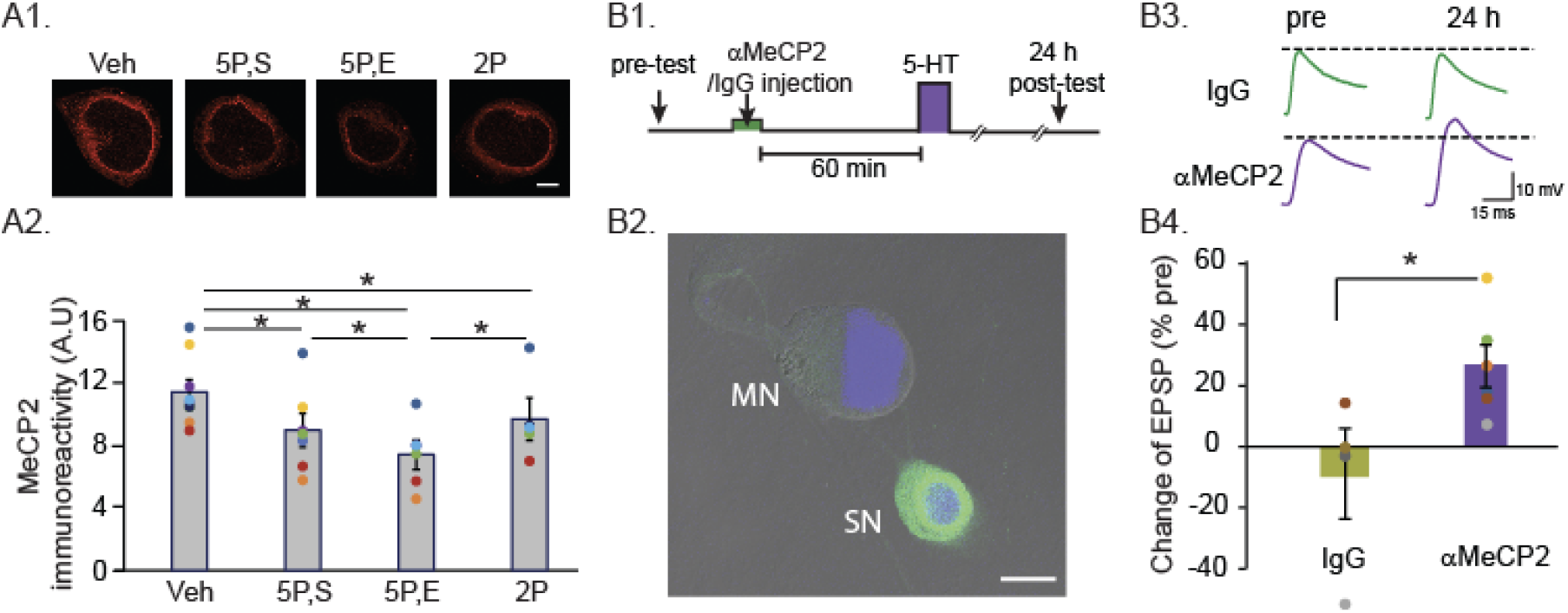
MeCP2 in the induction of LTF. A: Down-regulation of MeCP2 by LTF-inducing protocols. Immunofluorescence measurements of MeCP2 in isolated SNs 2 h after the Standard protocol (5P,S), Enhanced protocol (5P,E) and two-pulse protocol (2P) vs. Veh. A1: Representative confocal images (scale bar, 20 μm). A2: Summary data. Two h after the end of repeated 5-HT treatment, MeCP2 was significantly decreased in the cell bodies (cytoplasm and nucleus) for all protocols. Moreover, the decrease in MeCP2 in 5P,E group was significantly different from 5P,S and 2P groups. There was no significant difference between 5P,S and 2P groups. B: One pulse of 5-HT paired with αMeCP2 induced LTF. B1: Protocol for control IgG or αMeCP2 injection and electrophysiological testing. Antibodies were injected in the cytosol of SNs after EPSP pre-test and 1 h before 5-HT treatment. B2: Differential interference contrast confocal imaging of a SN-MN co-culture after injection. DAPI (blue) was used to stain nuclei. Injections were deemed successful if the SN cytoplasm and processes were filled with fluorescein (green dye). Scale bar, 50 μm. B3: Representative EPSPs before (Pre) and 24 h after 1 pulse of 5-HT. B4: Summary data. αMeCP2 together with one pulse of 5-HT induced LTF at 24 h.

The observation that MeCP2 was reduced during the induction phase of LTF suggests MeCP2 may act as a transcription repressor, with relief of repression necessary to allow LTF formation. To test this hypothesis, we injected αMeCP2 into the cytoplasm of sensory neurons (SNs) in SN - motor neuron co-cultures, to remove the presumed repression. We then applied one 5-min pulse of 5-HT, a protocol that usually only induces short-term facilitation (STF) lasting for ∼10 min ^16, 17^ and examined whether blocking MeCP2 function suffices to convert STF to LTF (Fig. 1B). Three groups were examined: 1) control IgG injection, treated with one pulse of 5-HT (IgG + 1×5-HT); 2) αMeCP2 injection, treated with vehicle (αMeCP2 + Veh); and 3) MeCP2 antibody injection, treated with one pulse (αMeCP2 + 1×5-HT). One 5-HT pulse induced a 28 ± 6% (n = 7) increase in EPSP amplitude at 24 h post treatment in the αMeCP2 + 1×5-HT group. One-way ANOVA indicated a significant difference among the groups (F_2,18_ = 7.75, p = 0.004). Subsequent pair-wise comparisons (SNK) indicated one pulse of 5-HT induced significant facilitation in the αMeCP2-injected co-cultures compared to IgG + 1×5-HT (n = 7, q = 5.387, p = 0.004) and αMeCP2 + Veh (n = 7, q = 3.915, p = 0.013) (Fig. 1B4), suggesting inhibition of MeCP2 activity facilitates induction of LTF. Injection of αMeCP2 alone did not significantly affect synaptic transmission at 24 h, compared to IgG + 1×5-HT (q = 1.472, p = 0.312). Importantly, there were no significant differences in basal synaptic strength (pre-test) (one-way ANOVA, F_2,18_ = 2.684, p = 0.95) and passive properties of the motor neurons (MNs) among the groups (resting potentials, F_2,18_ = 1.752, p = 0.202; input resistances, F_2,18_ = 1.155, p = 0.337) at 24 h post treatment. Therefore, the observed differences in facilitation between αMeCP2- and control IgG-injected co-cultures were not due to differences in basal synaptic strength or changes in these biophysical properties of the MNs. In addition, MeCP2 had no effect on the induction of STF. Immediately after exposure to one pulse of 5-HT, there was no significant difference between the facilitation in IgG-injected group (51 ± 14%, n = 7) and that in αMeCP2-injected group (57 ± 17%, n = 9) (Student t-test, t_14_= 0.269, p = 0.792).

Finally, we examined the roles of MeCP2 in the maintenance of LTF. As shown in Fig. 2, LTF induced by the Standard protocol disappears by 48 h relative to pre-test (93 ± 8.9 %, n = 7) and also is not present at 5 d (73 ± 4.4 %, n = 6). However, in the αMeCP2-injected group, the increases in EPSP amplitude induced by the Standard protocol persisted to 48 h (124 ± 8.1 %, n = 7) and also at 5 d (134± 20.5 %, n = 6) (Fig. 2C). In addition, at 24 h post 5-HT, the EPSP increase with αMeCP2 was 147 ± 12.7 % of pre-test (n = 7), as compared to 127 ± 9.5 % (n = 8) with IgG. Statistical analyses (two-way repeated measurement ANOVA) indicated significant overall differences in the amplitude of EPSPs among the IgG- and MeCP2-injected groups (*F*_*1,22*_ = 16.60, p < 0.001), and also among the three time points (*F*_*2,22*_ = 4.98, p = 0.016). These results indicate αMeCP2 can prolong the duration of LTF induced by the Standard protocol, suggesting MeCP2 may contribute to the maintenance of LTF.

**Figure 2.**
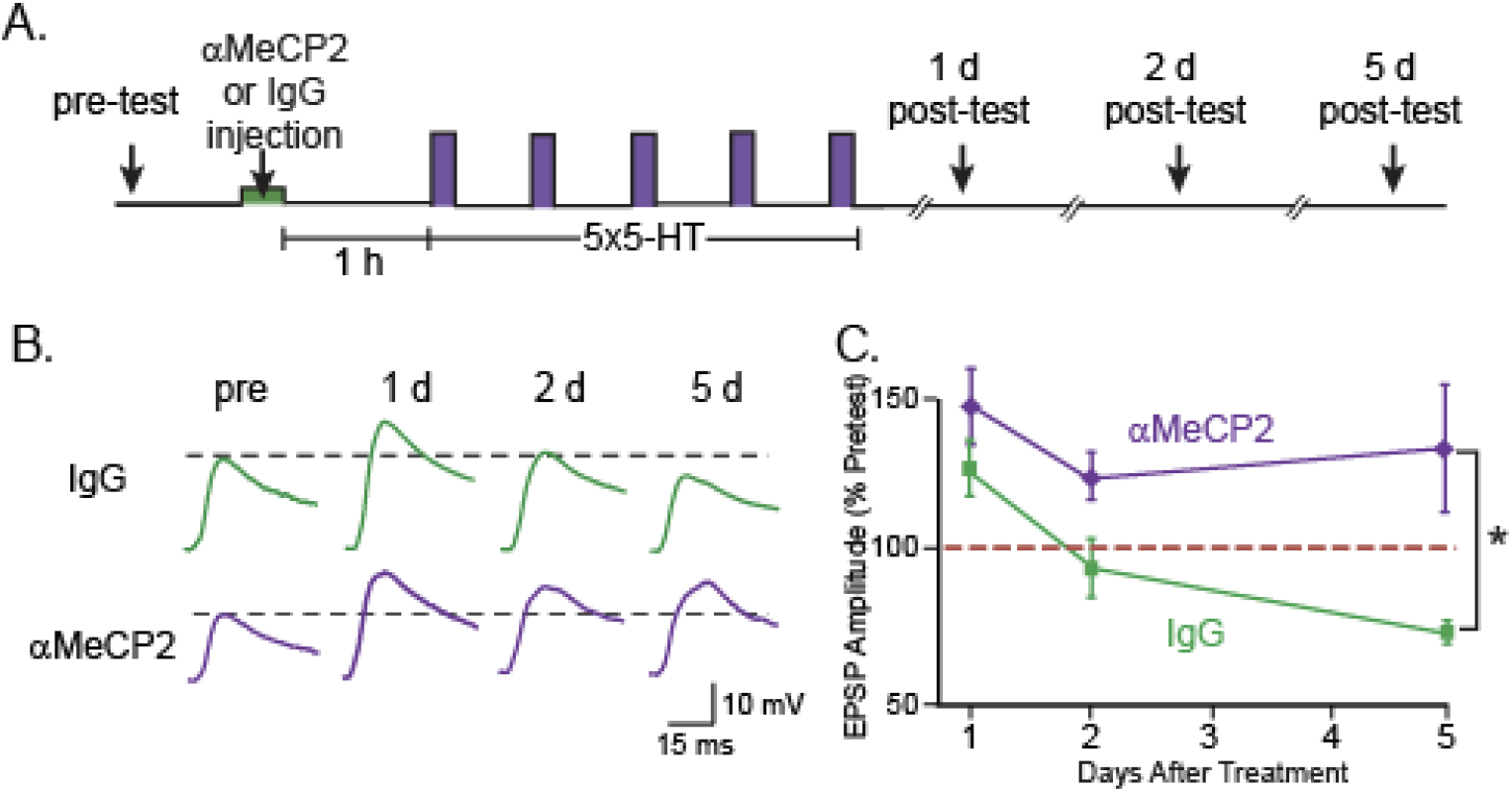
MeCP2 in the maintenance of LTF. A: Protocol for control IgG or αMeCP2 injection and electrophysiological testing. Antibodies were injected in the cytosol of SNs after the EPSP pre-test and 1 h before treatment with the Standard protocol. B: Representative EPSPs before (Pre), 1, 2 and 5 d after 5 pulse of 5-HT. C: Summary data. Injection of αMeCP2 together with the Standard protocol induced LTF lasting at least 5 days.

Both total and phosphorylated CREB1 are upregulated to activate gene transcription required for LTF 2 h after 5-HT treatment^18^, at the same time point we observed reduced MeCP2 levels (Fig. 1A). This decrease in MeCP2 is consistent with previous findings that MeCP2 levels decrease in cultured cortical neurons in respond to treatments that lead to a potentiation of neuronal response^19^ and in response to forskolin and KCl treatments in postnatal day 1 cortical neurons^20^. These results suggest MeCP2 may function as an inhibitor of synaptic plasticity. One pulse of 5-HT, which normally induces only STF, when paired with injection of αMeCP2 into SNs to remove presumed repression of transcription by MeCP2, sufficed to induce LTF (Fig. 1B), confirming MeCP2 may act as an inhibitory constraint on LTF. Interestingly, injection of αMeCP2 into SNs had no effect on basal synaptic transmission 24 h after injection. This finding is in contrast to studies that report changes in basal synaptic transmission^7, 10-12, 19, 21-23^. This difference may be due to the acute effects of antibody injection *vs*. long-term reductions of MeCP2 levels in mutant mice, with the latter possibly having substantial presynaptic and postsynaptic effects in neural circuits.

The most striking finding is that removing the inhibitory effects of MeCP2 by αMeCP2 injection prolongs LTF. These data provide initial evidence for a role of MeCP2 in memory maintenance. With the antibody, LTF induced by the Standard protocol persisted at least 5 d. Previously this persistence necessitated either the Enhanced protocol^15^, or two repetitions of the Standard protocol on consecutive days^24^.

CREB2 is another transcription repressor important for the induction of LTF, and 5-HT leads to relief of CREB2-mediated repression^16^. Interestingly, the regulation of MeCP2 and CREB2 appears to display different dynamics, but they have similar effects on LTF. The functional significance of these distinct dynamics requires further investigation.

A simplified model that incorporates the suggested role of MeCP2 in the induction of LTF is illustrated in Fig. 3. 5-HT triggers several kinase cascades to regulate transcription factors including: 1) PKA-dependent activation of CREB1, which initiates transcription of genes essential for LTF, such as *c/ebp*^1*3*^; 2) ERK-dependent inactivation of CREB2, relieving transcriptional repression by CREB2^16^; 3) convergence of PKA and ERK to activate p90 ribosomal S6 kinase (RSK), which activates CREB1^25^; and 4) ERK-dependent activation of the transcription activator C/EBP^26^. We hypothesize the 5-HT-induced decrease in MeCP2 expression is likely mediated in part by the CREB-dependent micro-RNA miR132^20^ (Fig. 3, dashed lines). Based on Bambah-Mukku et al.^27^, we further posit MeCP2 acts, at least in part, via interfering with binding of C/EBP to the promoters of effector genes required for LTF (Fig. 3).

**Figure 3.**
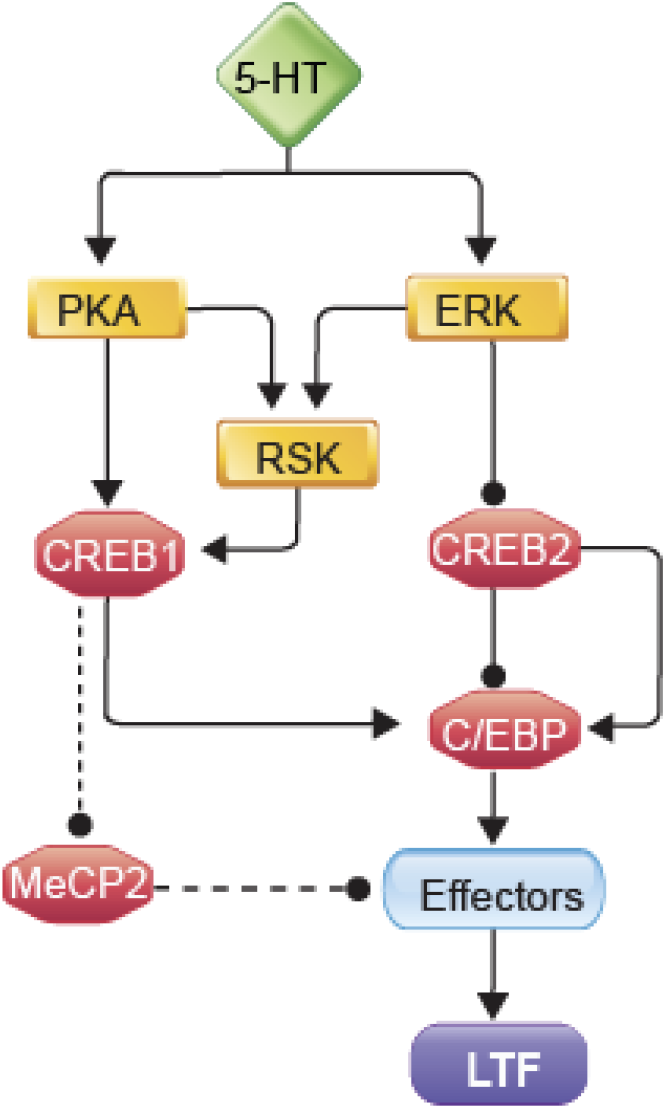
A simplified model incorporating MeCP2 into molecular pathways in SNs that regulate induction and maintenance of LTF. 5-HT initiates cascades that activate the key kinases PKA, ERK, and RSK. These kinases in turn activate CREB1 and C/EBP, and inactivate CREB2. 5-HT also down-regulates MeCP2, likely via CREB1, releasing repression of transcription and augmenting LTF (dashed line). Dashed lines represent multiple, mostly uncharacterized processes, which might occur sequentially or, in some cases, partly in parallel. Arrowheads, activation; circular ends, repression.

These results suggest that the *Aplysia* LTF model may serve as a powerful system to gain further insights into the roles of MeCP2, and plausibly other transcription modulators, in learning and memory, possibly providing new insights into the mechanisms of cognitive deficits associated with Rett syndrome. It will be important to further characterize the role of MeCP2 in synaptic plasticity by analyzing the dynamics of its expression at various times after different training protocols. It will also be important to investigate roles of MeCP2 phosphorylation, because phosphorylation of MeCP2 is involved in modulating LTP and memory^8^.

## Acknowledgements

We wish to thank E. Kartikaningrum and Ling Zhong for preparing the cultures, and C. Neveu for comments on the manuscript and illustration. This research was supported by NIH grant NS019895.

## Online Methods

Isolated sensory neurons (SNs) or SN - L7 motor neuron (MN) co-cultures from *Aplysia californica* (NIH *Aplysia* resource facility, University of Miami, Miami, FL) were prepared according to conventional procedures ^15, 25^. Three 5-HT protocols were used to induce LTF. These are: 1) the Standard protocol, five 5-min pulses of 50 μM 5-HT (Sigma, H9523) with interstimulus interval (ISI) of 20 min. Subsequently denoted “5P,S”, this protocol has been commonly used for decades^15, 28^; 2) the Enhanced protocol (5 pulses of 5-HT with computationally designed irregular ISIs as 10-10-5-30, denoted “5P,E”) which induces stronger LTF persisting at least 5 d^15^; and 3) the 2-pulse protocol denoted “2P”, with an ISI of 45 min, which induces LTF comparable to the Standard protocol^29^. A separate group of control cultures were treated with vehicle alone instead of 5-HT (Veh) (L15:ASW) with ISIs of the Standard protocol. Two hours after the end of treatment, anti-MeCP2 antibody (αMeCP2) was used to examine changes in MeCP2 levels in western blot and immunofluorescence analysis.

For western blots to validate αMeCP2, pleural ganglia were lysed in ice-cold buffer containing (in mM): 10 EDTA, 20 Tris (pH 7.5), 1 Na orthovanadate, 1 DTT, 2 NaF, 2 NaPPi, 0.5 okadaic acid, 1 PMSF, 1% SDS, 1% protease inhibitor cocktail (Sigma, St. Louis, MO) and stored at -20 °C until further use. Thirty μg of *Aplysia* pleural extract were resolved using SDS-PAGE after protein amount was estimated by a modified Lowry method (DC protein assay, Bio-Rad, Hercules, CA). Approximately 20 ug protein extract from each sample was run on mini-protean TGX gel (Bio-Rad) and transferred to nitrocellulose membrane (Bio-Rad). Membranes were blocked in 5% milk and TBS with 0.2% Tween20 (TBST) overnight at 4 °C, and were then incubated with αMeCP2 overnight at 4°C (αMeCP2; Fisher, cat# PA1888) (1:1000). The membranes were then washed with TBST and incubated with a secondary antibody (Goat anti-Rabbit IgG (H+L) Cross-Adsorbed Secondary Antibody, HRP, Invitrogen, cat#31462) (1:1,000) for 2 h at room temperature. Immunoreactive bands were visualized by Amersham™ ECL™ Western Blotting Detection Reagents (GE Healthcare) and the membranes were scanned with SRX-101A film processor. Band intensity was analyzed with ImageJ-win64 software (NIH).

Preabsorption of αMeCP2 with a synthetic peptide, using the sequence derived from amino acids 1-15 of mouse MeCP2 protein (ThermoFisher, Cat# PEP-120), was conducted per standard procedure (Peptide: antibody 5:1, incubated for 24 h at 4 °C)^30^.

Immunofluorescence procedures followed those used in previous studies^31^. Briefly, cells were fixed in a solution of 4 % paraformaldehyde in PBS containing 30% sucrose^32^. After three rinses in PBS, fixed cells were blocked for 30 min at room temperature in Superblock blocking buffer (Pierce, Holmdel, NJ) / 0.2 % Triton X-100 / 3 % normal goat serum and subsequently incubated overnight at 4 °C with αMeCP2 (1:500) diluted in blocking solution. Secondary antibody (goat anti-rabbit secondary antibody conjugated to Cy-3, Jackson Lab, 1:200 dilution) was applied in the same blocking solution for 1 h at room temperature. Cells were then mounted using Prolong anti-fade medium (Invitrogen, Carlsbad, California). Images were obtained with Zeiss LSM800 confocal microscope using a 63× oil immersion lens. A z-series of optical sections through the cell body (0.5 μm increments) was taken, and the section through the middle of the nucleus was used for analysis of mean fluorescence intensity with ImageJ.

Five to 10 neurons on each coverslip were analyzed, and measurements from neurons on the same coverslip were averaged. These experiments were performed in a blind manner so that the investigator analyzing the images was unaware of the treatment the SNs received. The number of samples (n) reported indicates numbers of dishes assessed.

For antibody injection, pressure injection of αMeCP2 (0.5 μg/μl) or rabbit IgG in injection buffer [100 mM KCl, 2.5 mg/ml 70 kD fluorescein-dextran (Invitrogen, Carlsbad, CA)] into sensory neuron cytoplasm was done before 5-HT treatment using the Eppendorf microinjection system. The efficiency of the injection was monitored with a fluorescence microscope. The experimenter performing the electrophysiological testing was “blind” to the identity of the antibody injected and to the treatment (vehicle *vs*. 5-HT). After the 24-h post-test, a subset of co-cultures injected with anti-MeCP2 or IgG antibody was fixed with 4% paraformaldehyde and counterstained with DAPI (4′, 6-diamidino-2-phenylindole, Jackson ImmunoResearch, West Grove, PA), in order for confocal imaging to visualize the intracellular localization of the injected primary antibody and localization of the MN and the SN (Fig. 1B).

For electrophysiological experiments, sensorimotor co-cultures were prepared as described previously ^32^. Excitatory postsynaptic potentials (EPSPs) were recorded from motor neurons with 10-20 MΩ sharp electrodes filled with 3 M potassium acetate. Neurons with resting membrane potential more positive than -40 mV and input resistance smaller than 10 MΩ were excluded. Stimulation of presynaptic SNs was performed extracellularly using a blunt patch electrode filled with modified L15 - ASW. Data acquisition was performed using pClamp (Molecular Devices).

Synaptic facilitation was defined as an increase in the EPSP amplitude in either the IgG-or αMeCP2-injected groups as compared to EPSPs measured in the pretest. For statistical analysis of LTF, the amplitude of the EPSP in mV was measured before (pre-test), 1 day (also 2 and 5 days for Fig. 2) after (post-test) exposure to 5-HT (50 μM) or vehicle, and the post/pre-ratio was examined. To examine STF, EPSPs were tested 1 h after αMeCP2 or IgG injection. After the initial EPSP (pretest), a bolus of 5-HT was added to the SN-MN co-cultures to achieve a final concentration at 50 μM. Five min later the EPSP was retested (post-test).

## Supplementary Information

**Figure S1:**
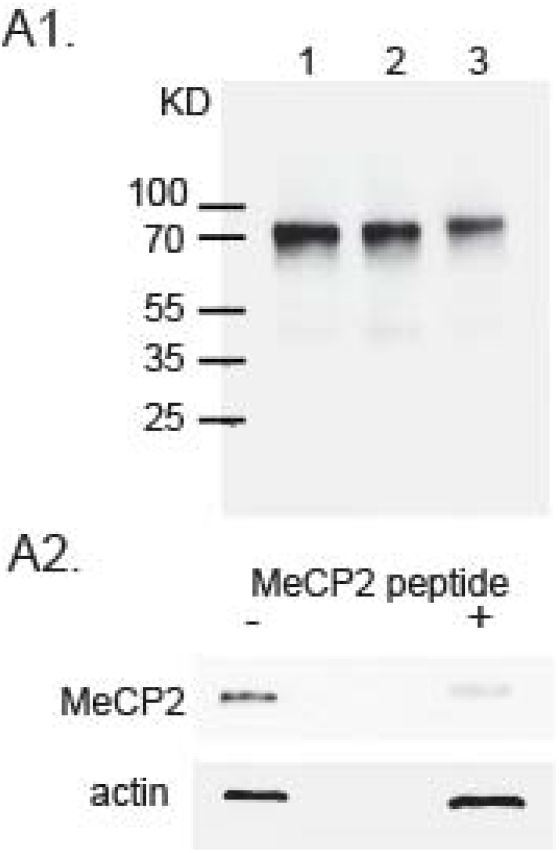
Validation of the specificity of αMeCP2 in *Aplysia*. A1: Western blot analysis indicated αMeCP2 recognized a single band with the expected molecular weight of 75 kD ^33-35^. Lanes 1–3 are lysates from the pleural ganglia of three animals. A2: Immunoreactivity to MeCP2 was diminished by αMeCP2 pre-absorption with a synthetic MeCP2 peptide. Actin served as a loading control.

